# *Incilius alvarius* cell-based synthesis of 5-MeO-DMT

**DOI:** 10.1101/2022.05.20.492789

**Authors:** Leonard Lerer, Eric Reynolds, Jeet Varia, Karin Blakolmer, Bernard Lerer

## Abstract

There is growing interest in the therapeutic potential of 5-MeO-DMT (5-methoxy-N,N-dimethyltryptamine) for psychiatric disorders. While 5-MeO-DMT can be chemically synthesized, the parotoid gland secretions of *Incilius alvarius* (also known as the Colorado River or Sonoran Desert toad) contain 5-MeO-DMT and other molecules including bufotenine, bufagenins, bufotoxins, and indole alkylamines that may have individual clinical utility or act as *entourage molecules* to enhance the activity of 5-MeO-DMT. *Incilius alvarius* is currently under severe ecological pressure due to demand for natural 5-MeO-DMT and habitat loss. We established a cell line from tissue obtained by wedge biopsy of the *Incilius alvarius* parotoid gland and confirmed the cell-based biosynthesis of 5-MeO-DMT by LC-MS/MS. Cell-based biosynthesis of *Incilius alvarius* parotoid gland secretions is a potentially cruelty-free and sustainable source of naturally derived 5-MeO-DMT for research and drug development.

**Graphical Abstract:** 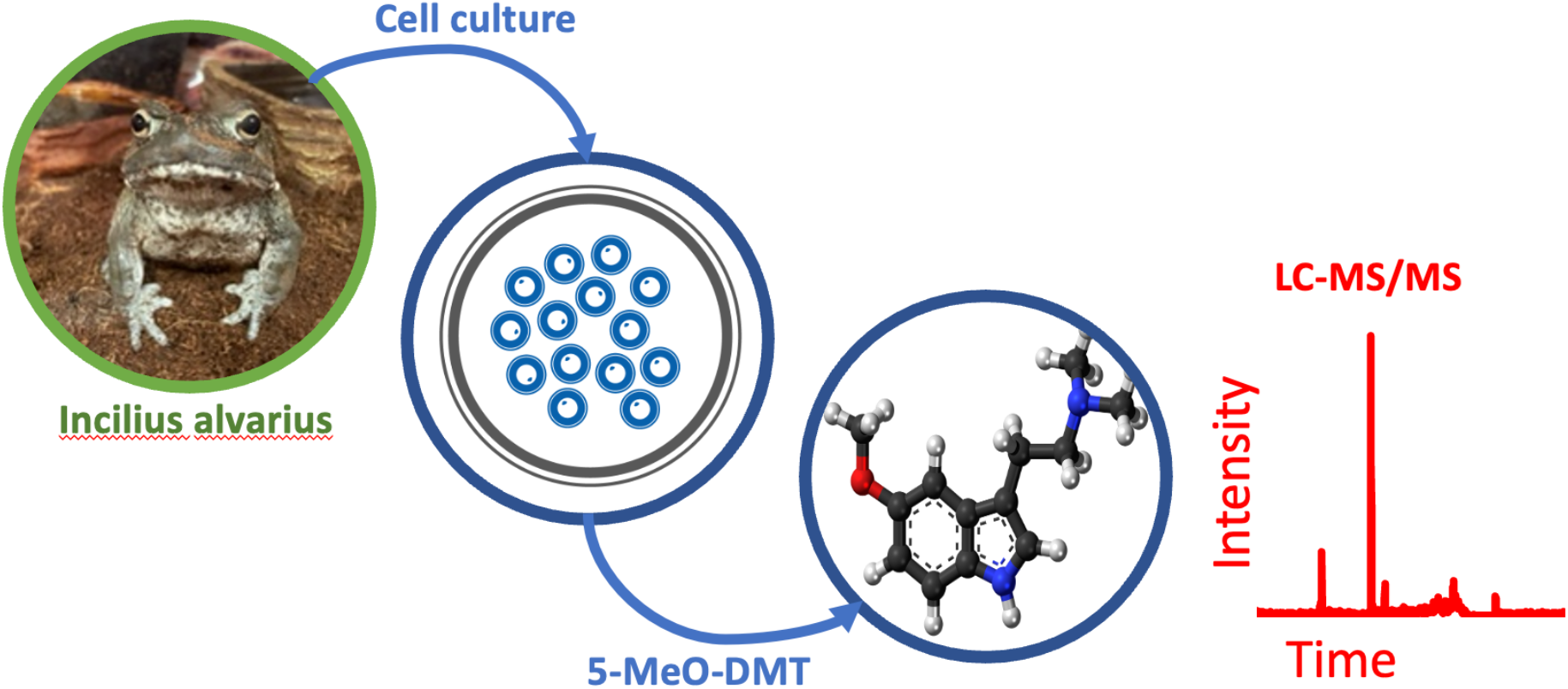

## 1. Introduction

5-methoxy-N,N-dimethyltryptamine (5-MeO-DMT) is the primary psychoactive component of the parotoid gland secretion (“venom”) of *Incilius alvarius,* the Sonoran Desert or Colorado River toad, and accounts for 20-30% of the dry weight of the secretion (Uthaug et al., 2019). 5-MeO-DMT primarily acts as an agonist at the 5-HT_1A_ and 5-HT_2A_ receptors with a higher affinity for the 5-HT_1A_ subtype (Ray, 2010; Halberstadt and Geyer, 2011). 5-MeO-DMT is O-demethylated by polymorphic cytochrome P450 2D6 (CYP2D6) to an active metabolite, bufotenine (Wieland et al., 1934; Erspamer et al., 1965; Erspamer et al., 1967).

Neuropsychiatric disorders including mood and anxiety disorders, are leading causes of disability worldwide and place an enormous economic burden on society (Gustavsson et al., 2011; Whiteford et al., 2013; NIH, 2022). Serotonergic psychedelics are receiving increasing attention as novel therapeutics for depression and other psychiatric and neurological disorders (Coppola et al., 2022). Preclinical and clinical evidence support neuroplasticity as the convergent, downstream mechanism of action of psychedelics. Through their primary glutamate or serotonin receptor targets, psychedelics including psilocybin, lysergic acid diethylamide (LSD), 5-MeO-DMT, and N,N-dimethyltryptamine (DMT) induce synaptic, structural, and functional changes, particularly in the pyramidal neurons of the prefrontal cortex (Olson, 2022). 5-MeO-DMT appears to be pharmacodynamically unique as compared to other psychedelics in terms of the intensity and rapid onset of action, short duration of effect, and transcriptomic and other parameters (Dakic et al., 2017). An optimized chemical method for the synthesis of 5-MeO-DMT has been described (Sherwood et al., 2020). However, volunteers consuming toad secretion containing 5-MeO-DMT experienced a 20–30% increase in the magnitude of subjective effects including “ego dissolution” and “altered states of consciousness” compared to the effects reported by volunteers who used synthetic 5-MeO-DMT (Shulgin and Shulgin, 1997; Shen et al., 2010; Uthaug et al., 2019).

It is possible that the non-psychedelic molecules in the toad secretion, also known as the *entourage molecules,* may be responsible for the enhanced effect of toad-derived 5-MeO-DMT, but whether or how the *entourage molecules* interact with the psychedelic molecules, thereby modulating the psychoactive experience and the neuroplastic effect, is currently unknown (Shulgin and Shulgin, 1997; Shen et al., 2010; Uthaug et al., 2019). While 5-MeO-DMT is present in several entheogenic plants, it should be noted that due to the high concentrations in the parotoid secretions of *Incilius alvarius,* the Sonoran Desert toad is currently under severe ecological pressure due to demand for recreational, self-medication, and spiritual use (Wake and Vredenburg, 2008).

## 2. Methods

### 2.1 Parotoid gland biopsy

*Incilius alvarius* parotoid anatomy and morphology were established according to O ‘donohoe et al. (O’ donohoe et al., 2019). A partial parotoid gland biopsy was undertaken under aseptic conditions in two anesthetized *Incilius Alvarius* toads in conformity with the current best practice for the handling of laboratory animals. A slightly modified procedure as described by Semoncelli et al. (Simoncelli et al., 2015) was performed; full-thickness ventral skin was removed, and glandular tissue was cut with a scalpel into approximately 1 cm^3^ sections. Explants were then rinsed with 70% isopropanol, and phosphate-buffered saline and stored in an ultra-cold freezer until culture initiation.

### 2.2 Cell culture, immortalization, and maintenance

Basal media preparation described by Ellinger et al. (Ellinger et al., 1983) was employed with modification and composed of 2X L-15 (GIBCO) with an addition of 10% fetal bovine serum. Media was then supplemented with antibiotics, sugars, hormones, amino acids, and ionic compounds. Two treatment groups were established; gelatin (Sigma-Aldrich, St Louis, MO, USA) was applied to two 24-well plates while no adherent was applied to two 24-well plates. Explants were subsequently thawed in a 65°C water bath, rinsed with amphibian Ringers solution prepared as described by Handler et al. (Handler et al., 1979), resuspended in fresh media, and prepared for connective tissue digestion. Two treatments of cell isolators were used, collagenase was employed similarly to Okumoto et al. (Okumoto, 2001) followed by ECDS (enzyme-free cell dissociation solution). Upon completion, dissociated cells were strained *via* both 0.40 and 0.22 *μ*m cell strainers. Isolated cells were aseptically seeded into 24-well plates and incubated at 25°C and 5% CO_2_. Upon confluence, cells were disassociated with the initial reagent used and passaged to additional well plates. Media exchange occurred approximately every 3-4 days and cell immortalization was achieved using the SV40 T Antigen Cell Immortalization Kit (Alstem, Richmond, CA, USA). Cell culture was undertaken in an incubator at 25° C and 5% CO_2_. Upon confluence, the cells were disassociated and passaged with media exchange every 3-4 days. The culture was maintained for 45 days.

### 2.3 LC-MS Analysis

Before analysis, cell media was filtered with an Amicon Ultra-0.5 Centrifugal Filter Unit (MWCO 10 kDa, Millipore Sigma, Burlington, MA, USA). 5-MeO-DMT was quantified using a UPLC – Xevo TQ-S micro MS system (Waters Corporation, Milford, MA, USA). The calibration curve was prepared using the 5-MeO-DMT analytical standard, in methanol, from Cerilliant Corporation (Round Rock, Texas, USA) using a concentration range of 1-2000 ng/mL. For quantification, monitored ions at 130.25 m/z and 159.07 m/z were used. The column used was a Phenomenex (Torrance, CA, USA) Luna Omega 3 *μ*m Polar C18 (150 mm × 4.6 mm) at 40 °C, solvent A was 0.1 % formic acid and solvent B was acetonitrile and 0.1 % formic acid at a flow rate of 0.6 mL/min. The injection volume is 1 uL. The MS was used in positive ion mode with time-scheduled multiple reaction monitoring (MRM) acquisition. The source temperature was 150 °C, capillary voltage 1.00 kV, desolvation temperature 600 °C, desolvation gas flow of 1000 L/hr, with a cone gas flow of 75 L/hr.

In the positive-ion mode and under turbo-ion-spray ionization conditions, 5-MeO-DMT gave a precursor ion (M+H)+ of m/z 219.20. Product ion of m/z 159.07 and 130.25 was found to be predominant for 5-MeO-DMT under the collision energy of 40 V (the MRM method was developed by utilizing IntelliStart Software, Waters Corporation, Milford, MA, USA). The MRM transitions m/z 219.2→159.07 and 219.2→130.25 were chosen to analyze 5-MeO-DMT, which offered the strongest signal compared to other MRM transitions.

## 3. Results and Discussion

Parotoid cells obtained from biopsies of anesthetized *Incilius alvarius* toads were successfully immortalized. Cultures were maintained for 45 days. Dried media from the cell culture had a light tan appearance that was similar to the known appearance of dried *Incilius alvarius* parotoid secretion. Well plate media samples were collected at 12 and 36 days from initiation and were analyzed for the presence of 5-MeO-DMT using LC-MS/MS and compared to spontaneously secreted material from an *Incilius alvarius* toad. MS fragmentation of the media samples and secreted material showed conformity in structure with 5-MeO-DMT (Figure 1). Additional chromatographic peaks were present but were not identified due to the paucity of the material.

**Figure 1.**
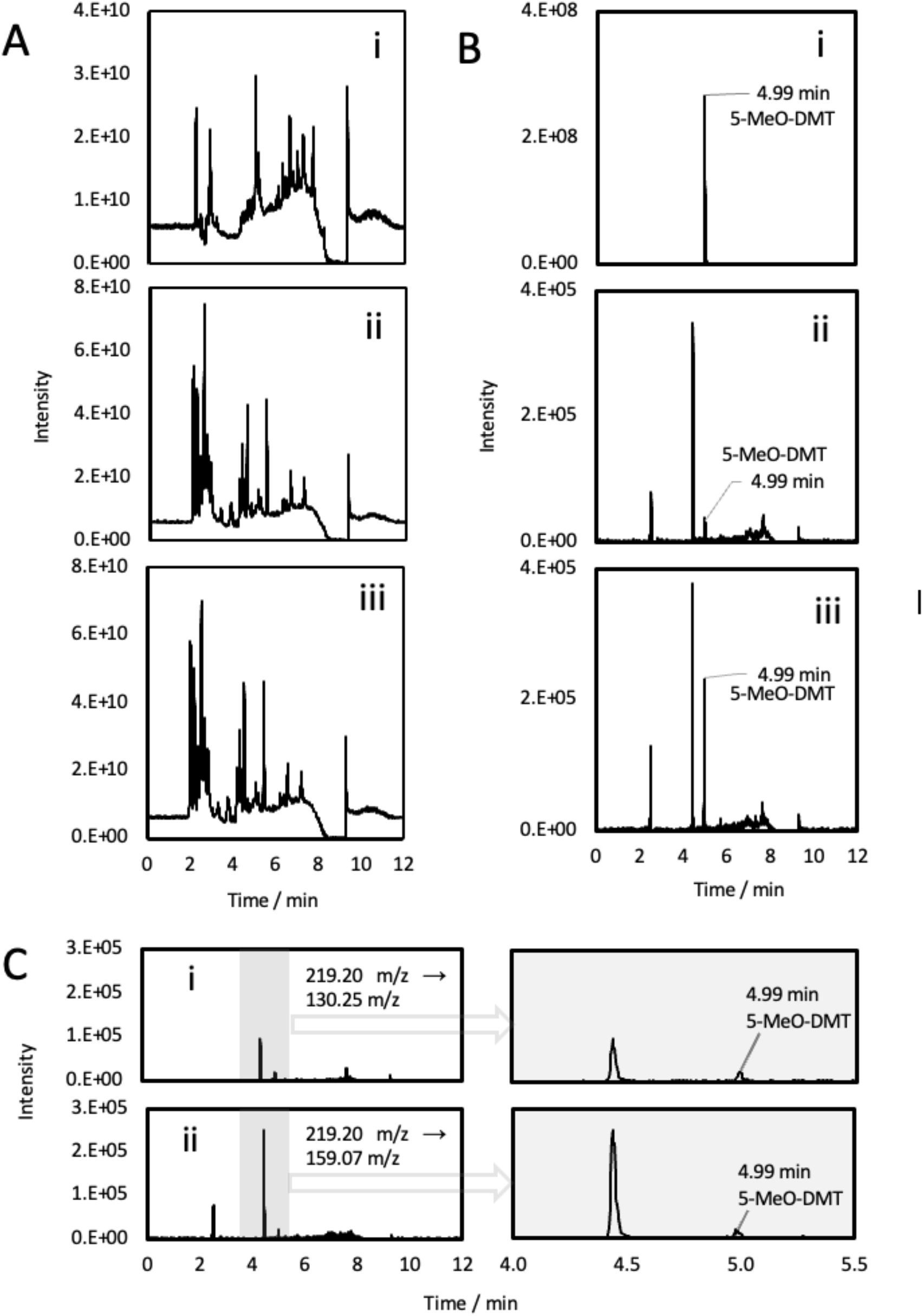
Chromatograms of 1 μL injection of (i) parotoid secretion (ii) parotoid cell-free media and (iii) blank media + 1 ng/mL 5-MeO-DMT. (A) Full scan MS, total ion chromatogram (TIC) (B) Sum of multiple reaction monitoring (MRM) transitions of 219.20 → 159.07, 219.20 → 130.25. (C) Identification and quantification of MRM transitions of 219.20 → 159.07 and 219.20 → 130.25.

In this pilot study, small quantities were produced, and it was not possible to accurately determine concentration or yield. We are conducting research to demonstrate scalable production of 5-MeO-DMT in the parotoid cell line and further confirm *de novo* biosynthesis. This work demonstrates the potential availability of naturally derived 5-MeO-DMT produced through cell culture, as opposed to the cruel and destructive practice of “milking” *Incilius alvarius* and supports efforts to ensure the protection of our planet’s entheogen heritage.

## Author Contributions

The manuscript was written with the contributions of all authors. All authors have approved the final version of the manuscript.

## Acknowledgements

The authors would like to thank Geovannie Ojeda-Torres for his chromatography work, Eric Kawka for comments and analysis of chromatograms, Dr. Sophia Gill, and Matthew Meifert for conducting the biopsies and postoperative follow-up, Jay Pleckham for general support and Katherine Spear for the diligent and loving care of the toads.

## Funding Acknowledgements

The authors received no financial support for the research, authorship, and/or publication of this article.

